# Craniofacial cartilage organoids from human embryonic stem cells via a neural crest cell intermediate

**DOI:** 10.1101/2021.05.31.446459

**Authors:** Lauren Foltz, Tyler Levy, Anthony Possemato, Mark Grimes

## Abstract

Severe birth defects or major injuries to the face require surgical reconstruction and rehabilitation. The ability to make bona fide craniofacial cartilage – cartilage of the head and face – from patient-derived induced pluripotent stem cells (iPSCs) to repair these birth defects and injuries has tremendous translational applications, but is not yet possible. The neural crest is the normal developmental pathway for craniofacial cartilage, however, the knowledge of cell signaling pathways that drive neural crest differentiation into craniofacial chondrocytes is limited. Here we describe a differentiation protocol that generated self-organizing craniofacial cartilage organoids from human embryonic stem cells (hESCs) and IPSCs through a neural crest stem cell (NCSC) intermediate. Histological staining of cartilage organoids revealed tissue architecture typical of hyaline cartilage. Organoids were composed of rounded aggregates of glassy, gray matrix that contained scattered small nuclei in lacunae. Mass spectrometry shows that the organoids express robust levels of cartilage markers including aggrecan, perlecan, proteoglycans, and many collagens. Organoids expressed markers indicative of neural crest lineage, as well as growth factors that are candidates for chondrocyte differentiation factors. The data suggest that chondrocyte differentiation is initiated by autocrine loops driven by a combination of secreted growth factors that bind to chondrocyte receptors. Craniofacial cartilage organoids were continuously cultured for one year, reaching up to one centimeter in diameter. The ability to grow craniofacial cartilage from NCSCs provides insights into the cell signaling mechanisms of differentiation into craniofacial cartilage, which lays the groundwork for understanding mechanistic origins of congenital craniofacial anomalies and repairing cartilaginous structures of the head and face.

## Introduction

Craniofacial reconstruction is necessary to treat several conditions including craniofacial birth defects, damage caused by head and neck cancer treatment, and traumatic facial injuries. These cases not only affect function but may also cause or contribute to psychological and social difficulties. It is difficult to reconstruct natural features with plastic surgery techniques, however, and transplanted tissue is often rejected by the recipient without immunosuppressants. The use of stem cells to grow craniofacial cartilage is an exciting prospect for effective regeneration and repair of birth defects and injuries to the head and face.

Craniofacial cartilage is comprised of hyaline cartilage for the nose and developing bones of the head, elastic cartilage for the larynx, epiglottis, and external ear, and fibrocartilage for temporomandibular joints. Unlike articular cartilage, craniofacial cartilage originates during development from chondrocytes that are derived from the neural crest. Neural crest cells (NCCs) are a developmental cell lineage restricted to vertebrates, and are sometimes considered the fourth germ layer. During neurulation, NCCs are specified at the neural plate border between the ectoderm and developing neuroepithelium. At the crest of the neural tube, NCCs undergo an epithelial to mesenchymal transition, delaminate, and migrate throughout the developing embryo (Bronner and LeDouarin, 2012). During this process, NCCs begin differentiating into a number of distinct cell types including cells of the peripheral nervous system, melanocytes, smooth muscle, and craniofacial cartilage and bone. Depending on their positioning and migratory patterns along the rostrocaudal axis of the developing embryo, NCCs can be further categorized into four main populations: cranial neural crest cells (CNCCs), cardiac neural crest cells (CaNCCs), vagal neural crest cells (VNCCs), and trunk neural crest cells (TNCCs). CNCCs eventually differentiate into craniofacial cartilage and bone, as well as glia, Schwann cells and cranial sensory glia (Bronner and LeDouarin, 2012). Therefore, craniofacial cartilage has a different developmental origin from the articular hyaline cartilage and fibrocartilage that composes the intervertebral disks and insertions of tendons in the rest of the body. In these tissues, the skeleton is first made of cartilage by chondrocytes derived from the mesoderm, which ossifies into bone; and some cartilage remains in selective locations, for example at the interface of joints (Chen et al., 2017).

Recent research efforts and clinical trials have focused on repairing articular cartilage defects. Several of these approaches use transplantation of mesenchymal stem cells and/or expanded chondrocytes or recruitment from subchondral bone (microfracturing) as a means to regenerate damaged joint and intervertebral cartilage (Burnsed et al., 2016; Guo et al., 2018; Huey et al., 2012). While current protocols can successfully generate functional cartilage, they suffer from a few major drawbacks. First, cartilage produced using embryoid body or mesoderm formation as a precursor has the potential to retain stem-like characteristics in the differentiated tissue, which could contribute to tumor formation (Fu et al., 2016; Maguire et al., 2015; Oldershaw et al., 2010; Soto et al., 2021). Second, the terminal differentiation stage of mesenchymal tissue is bone, and these systems can suffer from cartilage hypertrophy and ossification (Dexheimer et al., 2016; Van de Walle et al., 2018). In fact, transplanted cartilage derived from a mesenchymal origin can express heterogeneous markers for cartilaginous, fibrous, and hypertrophic tissues simultaneously (Huey et al., 2012; Steck et al., 2009). Finally, cartilage designed for joint repair has a different function from craniofacial cartilage and has to withstand applied stress (Choi et al., 2018; Huey et al., 2012; Panadero et al., 2016). In addition, microfracturing techniques are not applicable for craniofacial cartilage that has no adjacent bone. Developing techniques to effectively generate craniofacial cartilage is essential, requires procedures different from those used for other types of cartilage, and therefore requires an understanding of the cellular processes that underlie craniofacial cartilage development.

The exact growth factor signaling pathways directing the differentiation of human CNCCs into craniofacial cartilage is currently unknown, although insights can be gained from animal models (Van Otterloo et al., 2016). In general, chondrogenesis is under the control of several major developmental signaling pathways including from WNT, BMP/ TGFβ, and FGF family members, among others (Green et al., 2015; Mishina and Snider, 2014; Zhong et al., 2015). In mice, bone morphogenetic protein 2 (BMP2) and BMP4 are expressed in the cranial neural crest derived mesenchyme during E10.5-13.5 and signaling via the BMPRIa receptor is essential for proper palate, tooth, and temporomandibular joint formation (Bennett et al., 1995; Graf et al., 2016; Gu et al., 2014; Li and Chen, 2012; Liu et al., 2005; Mimura et al., 2016). Accordingly, mice lacking functional BMP receptors in the neural crest lineage exhibit hypoplastic mandibles, cleft palates, and die shortly after birth (Dudas et al., 2004; Komatsu et al., 2006). BMP7 has been shown to be able to induce extracellular matrix synthesis, prevent chondrocyte hypertrophy and maintain chrondrogenic potential (Mariani et al., 2014). BMP signaling crosstalk with the p53 apoptotic pathway controls nasal cartilage formation and fusion of the nasal septum in mice (Hayano et al., 2015).

Transforming growth factor β (TGFβ) family members have also been implicated in chondrogenesis, although the impact of different ligands to the formation of cartilage seems to be variable (Chimal-Monroy and Díaz de León, 1997; Yang et al., 2009). In particular, TGFβ-I and TGFβ-III have been used to increase chondrogenesis during in vitro differentiation of mesenchymal stem cells. Pre-seeding of TGFβ ligands on scaffolding materials is being investigated as a method to increase the mechanical strength of transplanted cartilage tissues (Deng et al., 2019; Pfeifer et al., 2019). These studies suggest that TGFβ ligands may be more important for cartilage derived from a mesenchymal stem cell origin.

Signaling pathways initiated by fibroblast growth factor (FGF) family members have also been shown to be important for both bone and cartilage formation in mice. Specifically, FGF8 has been shown to promote chondrogenesis over osteogenesis, both in the mouse skull and during thickening of the palate (Xu et al., 2018), and FGF18 has been known to induce cell proliferation and extracellular matrix synthesis in porcine and human cartilage (Mariani et al., 2014). In summary, most research examining growth factor signaling dynamics during craniofacial cartilage generation have been conducted using animal models and seems to have converged on BMP, TGFβ, and FGF signaling as the main signaling pathways involved in this process. However, exactly how each of these pathways conduct craniofacial development in human models is an area of active research.

Here we describe the generation of craniofacial cartilage organoids from human embryonic stem cells via a neural crest cell intermediate. We report a protocol for neural crest differentiation that integrates and streamlines several existing protocols from the recent literature. Using this protocol, craniofacial cartilage organoids were generated that did not require a scaffold, were amenable to handling and histological techniques, and did not rely on embedding in extracellular matrices. Tandem mass tag mass spectrometry and immunofluorescence revealed that craniofacial cartilage organoids expressed several markers of hyaline cartilage and retained markers indicative of their neural crest origin. We hypothesized that cartilage formation was triggered by an autocrine signaling loop. Using a subset of growth factors identified via mass spectrometry, we were able to accelerate organoid formation, suggesting that these pathways may play an important role in normal chondrocyte differentiation. Craniofacial cartilage organoids are a promising new model for this aspect of human development and may give insights into the signaling pathways that regulate cartilage formation.

## Results

### Neural crest stem cells differentiated into cartilage organoids

To differentiate craniofacial cartilage, we first directed the differentiation of human embryonic stem cells (hESCs) into a neural crest stem cell (NCSC) intermediate (Figure 1A). To do this we integrated a number of previously published protocols with the goal of shortening this stage of differentiation and generating the NCSC intermediate as quickly as possible while optimizing the yield. To obtain neural crest cells, it is first necessary to induce neuroectoderm formation. To do this, we used the LSB short protocol previously published by Kreitzer et al., which utilizes dual SMAD inhibition via the small molecules LDN193189 and SB431542, inhibiting BMP and TGF-β signaling respectively (Kreitzer et al., 2013a). However, we fit the timeline of this protocol to the research by Mica et al., which showed that early removal of TGF-β inhibitors, followed by the prolonged addition of the small molecule WNT signaling activator CHIR99021 (GSK3β inhibitor) increased anterior NCSC yield (Chambers et al., 2015; Mica et al., 2013). Finally, we added Shh, FGF8, ascorbic acid, and BDNF to the last stage of NCSC induction, following the neural crest specification protocol outlined by Zeltner et al., to further increase the number of NCSCs over a short time period (Zeltner et al., 2014). After neural crest differentiation, following the re-plating stages of the Zeltner et al. protocol, we then sorted NCSCs via magnetic-activated cell sorting based on p75/NGFR expression and re-plated cells as a monolayer on laminin and fibronectin substrate. Over the course of NCSC differentiation, hESCs acquired NCSC morphology and migrated outwards from hESC colonies (Figure 1B, Day 1-9).

**Figure 1.**
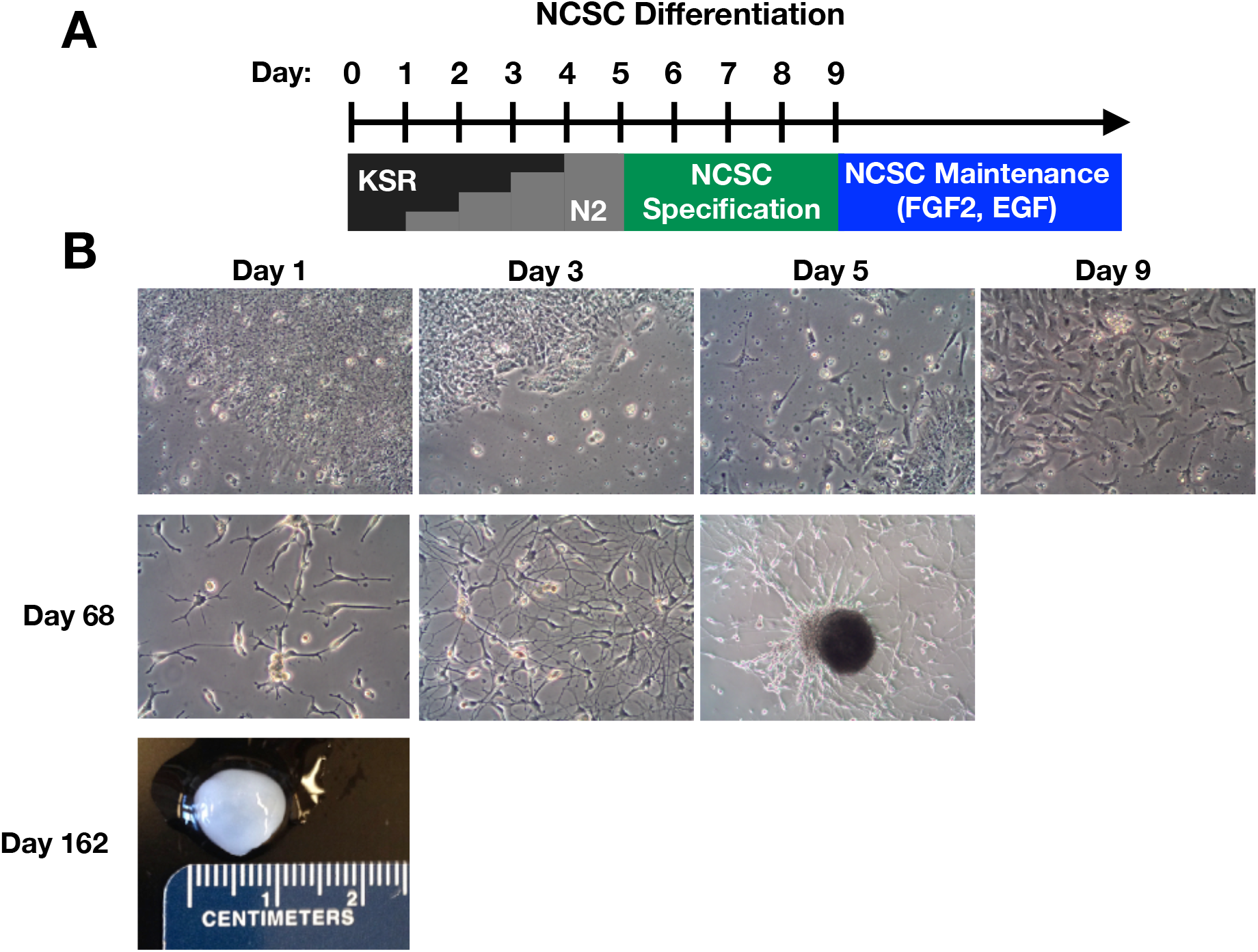
Neural crest differentiation protocol and cartilage organoids. **A)** Neural Crest Stem Cell (NCSC) differentiation protocol. WA09 hESCs were differentiated using stepwise replacement of KSR media with N2 for 5 days, followed by addition of NCSC specification media containing BDNF, FGF8, SHH, and ROCKi. Small molecule inhibitors were added to guide differentiation. NCSCs and organoid producing cells were maintained in N2 media containing FGF2 and EGF. **B)** Representative light microscopy images of cell morphology throughout differentiation and after prolonged culture. Day = number of days after the original differentiation protocol and does not represent the age of the pictured organoids. Organoids continue to spontaneously form in cell culture dishes containing differentiated NCSCs.

When left undisturbed for over 52 hours, confluent NCSCs migrated together and self-organized into roughly spherical growths on the surface of dishes (Figure 1B). Once formed, these organoids detached from the dish surface and floated in the cell culture media. Over time, cells remaining on the surface of the dish exhibited the morphology of migrating chondrocyte precursors (Figure 1B, Day 68). Floating organoids were easily harvested with a spatula, and remaining cells on the dish surface continued to produce new organoids. Cell confluency typically remained around 50% at this stage, with localized dense patches of cells that begin to migrate into new growths. The largest harvested organoid was one centimeter in diameter (Figure 1B, Day 162), and it was possible to maintain organoid-producing culture dishes for up to one year. Organoids had the appearance and physical characteristics of cartilage, being opaque and glassy in appearance, robust to handling, and resistant to applied pressure by springing back into shape.

### Organoids were positive for neural crest cell markers and were composed of collagens and other extracellular matrix proteins

To test if organoids generated from neural crest cells were cartilaginous, we first looked for markers of cartilage and the neural crest by immunofluorescence and staining of organoid cryosections by histological analysis and sectioning techniques. Hematoxylin and eosin staining of sections highlighted the typical morphology of cartilaginous tissues – small cell nuclei in the center of lacunae embedded in a glassy gray matrix (Figure 2A). The indicative basophilic staining is likely due to the presence of negatively charged aggrecan and other proteoglycans, which was confirmed by mass spectrometry (Figure 3). Staining with Saffranin O and Toluidine Blue was performed to verify collagen content and visualize collagen distribution throughout slices (Figure 2A). Immunofluorescence staining revealed that some cells in the center of organoid sections retained stem-like markers such as Sox2 (Figure 2B), although these cells were isolated and dispersed throughout sections. Neural cell adhesion molecule (NCAM) staining, a marker for immature neurons and early neural crest lineage, was also seen in the interior of larger organoids, although these cells appeared to have undergone apoptosis as evidenced by fragmented nuclei. Some cell death in the center of larger organoids was expected, we suspect that these organoids could only get so large without the use of a bioreactor due to the avascular nature of normal cartilage tissues. Doublecortin (DCX), a migratory neural crest and chondrocyte cell marker (Ge et al., 2014; Vermillion et al., 2014; Zhang et al., 2007), showed robust staining, particularly in the outer layer of organoids, suggesting outward growth (Figure 2B). Finally, COL2A1 staining, the primary collagen isoform found in craniofacial cartilage, exhibited bright and diffuse staining throughout organoids (Figure 2B). In summary, both immunofluorescence and histological staining methods indicated the presence of abundant collagen in organoids, consistent with formation of craniofacial cartilage. In addition, organoids retained some markers of their neural crest cell lineage.

**Figure 2.**
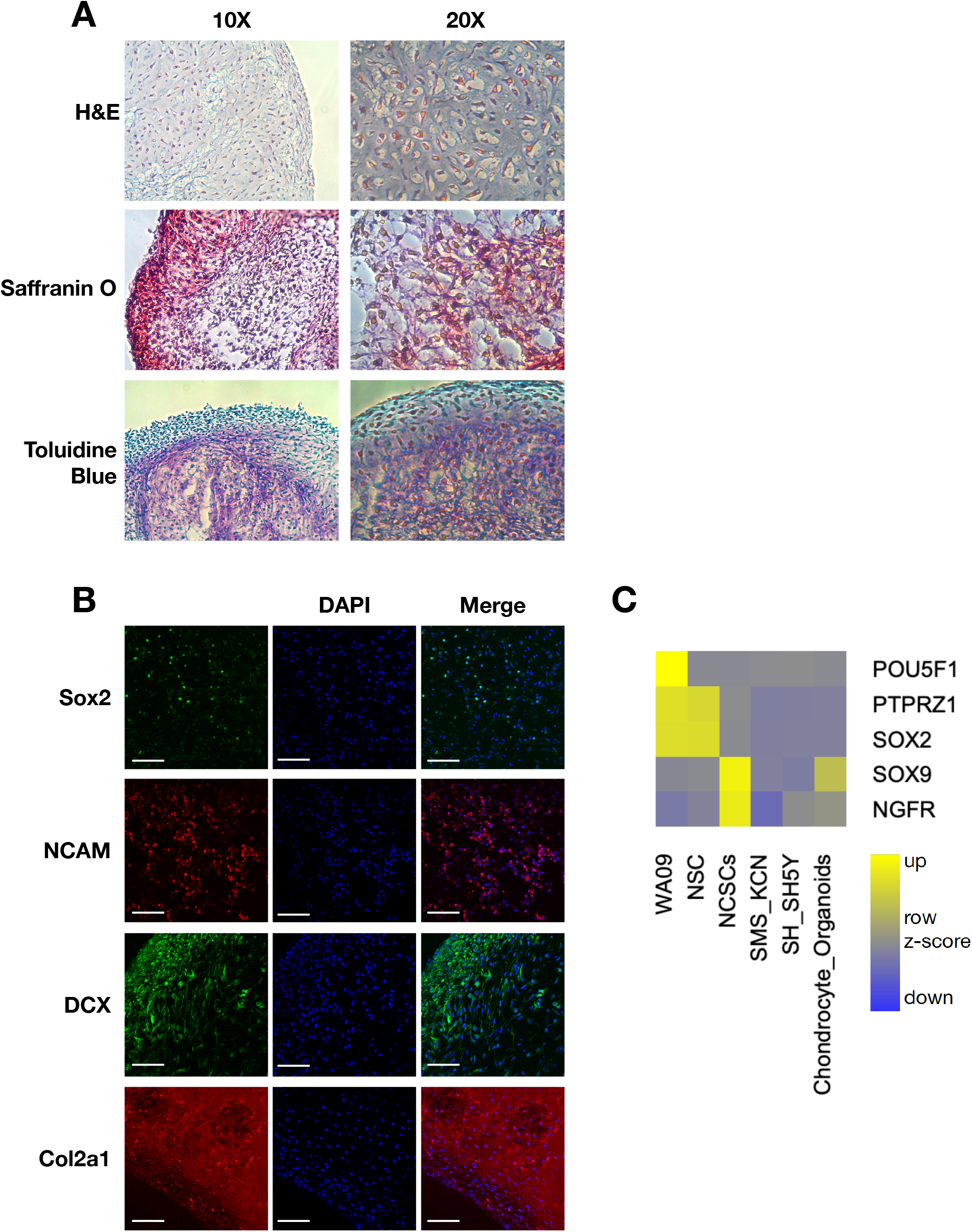
Organoid structure and markers for collagen and immature neurons. (A) Histology and simple stains of organoid cryosections. H&E, hematoxylin and eosin. Saffranin O stains collagen red. Toluidine Blue stains collagen purple. (B) Immunofluorescence of organoid cryosections. Sox2, pluripotency marker. NCAM, neural cell adhesion molecule. DCX, doublecortin, immature neuronal marker. Col2a1, collagen II. Scale bar = 100μm.

**Figure 3.**
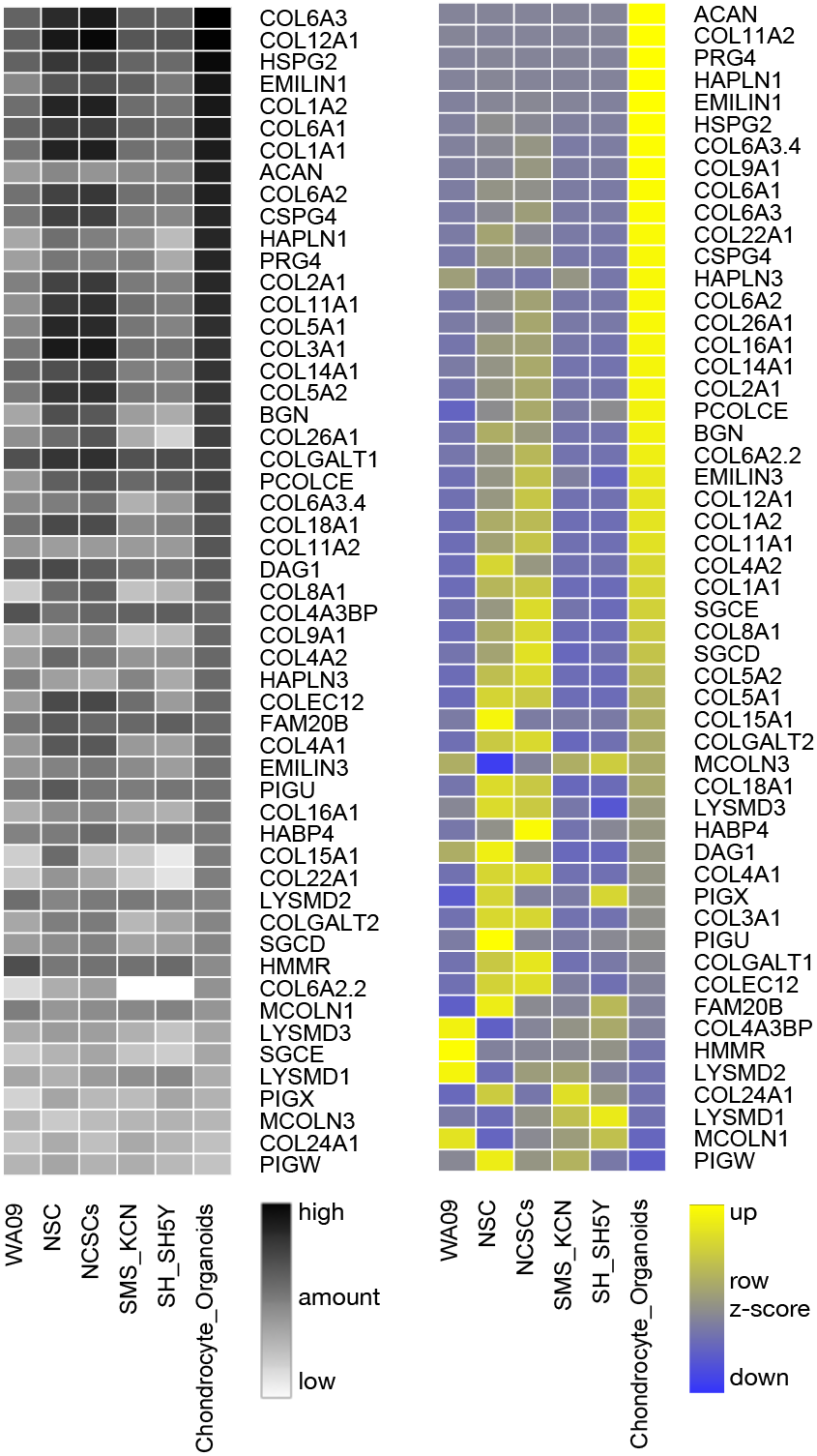
Summary of mass spectrometry identification of chondrocyte markers, plotted as greyscale heatmaps (left) of detected signals and row z-scores (right). All of the major cartilage proteins (*e*.*g*., ACAN, PRG4, and COL2A1) are expressed in high amounts. Also detected were perlecan and collagen 6 family members, which suggests that the cells are starting to make a pericellular matrix. TMT data were obtained in collaboration with Bluefin Biomedicine (Beverly, MA) comparing H9 human embryonic stem cells (WA09); neural stem cells (NSCs); neural crest stem cells (NCSCs); and neuroblastoma cell lines (SH-SY5Y, SMS-KCN) to chondrocyte organoids (rightmost column). Greyscale heatmaps (left) show relative amounts (log2 scaled) of proteins with black representing the maximum signal and white representing no signal. Color heatmaps (right) show data transformed by calculating the z-score of protein signals across samples (row z-score), with increased expression represented in yellow and decreased expression in blue. Data are ordered from highest to lowest by the chondrocyte organoid column.

To further investigate the composition of organoids, and the change in protein expression throughout the differentiation procedure, we performed tandem mass tag mass spectrometry (TMT-MS) analysis comparing the parent stem cell line (H9/WA09), neural stem cells (p75 negative population after magnetic sorting, NSCs), the p75 positive NCSCs, cartilage organoids, and two neuroblastoma cell lines (SMS-KCN and SH-SY5Y). Our TMT-MS approach enabled us to directly compare relative protein expression in all samples and therefore track changes at each stage of differentiation. In particular, this analysis reliably yields robust quantitative comparisons of proteins in different samples. The parent hESC line, WA09, expressed the highest amounts of the pluripotency markers OCT4 (POU5F1 gene) and SOX2 (Figure 2C). NSCs expressed high amounts of SOX2 protein, but significantly less OCT4 than undifferentiated stem cells. Compared to NSCs, NCSCs had higher amounts of the neural crest markers SOX9 and the low-affinity nerve growth factor receptor (NGFR), which was used to sort this cell population. Craniofacial cartilage organoid samples retained markers indicative of their neural crest origin (SOX9), and had low expression of OCT4 and SOX2 (Figure 2C).

We found that twenty chondrocyte markers in cartilage organoids are significantly elevated compared to NCSCs (p < 2.2 × 10^−16^, Welch two-sample t-test). Organoid samples were highly enriched for most major collagen isoforms, especially COL1A1 and COL2A1 (Figure 3). Aggrecan (ACAN), HAPLN1, HAPLN3, and EMILIN1 were also enriched, all of which are involved in cell-matrix attachments and hyaluronan and proteoglycan binding (Chen and Birk, 2013; Spicer et al., 2003). Aggrecan in particular is a major protein required for the structural integrity and durability of both hyaline and articular cartilage (Gibson and Briggs, 2016). Collagen X (COL10A1) and MMP13, markers for chondrocyte hypertrophy (Miao et al., 2018), were not detected. These data strongly reinforce the conclusion that organoids derived from the neural crest cell intermediate were craniofacial cartilage, one of the terminal differentiation stages of the neural crest lineage.

### Craniofacial cartilage organoids expressed ligand and receptor pairs that directed differentiation through the formation of an autocrine loop

In sum, 9372 proteins were identified in our TMT-MS panel, including 44 different growth factors and 185 cell signaling receptors. Because the signaling pathways involved in human craniofacial development have not been fully described, this data revealed which signaling pathways were represented at each stage of development. Receptor and ligand pairs that are known to be involved in either neural crest formation or cartilage differentiation were present in the dataset. Notably, enriched growth factors included MDK; PTN; TGF-β family members; WNT ligands; FGF1 and FGF2; HDGF; and VGF (Figure 4). Both PTN and MDK are important for chondrocyte differentiation and cartilage formation because they bind extracellular heparan-sulfate proteoglycans and cell surface syndecans as well as the signaling proteins ALK and PTPRZ1 (Pufe et al., 2007). In animal models and human cell lines, PTN and MDK have been shown to increase chondrocyte proliferation and both have been implicated in bone and cartilage repair after injury (Bouderlique et al., 2014; Ohta et al., 1999; Zhang et al., 2010). Several growth factor receptors were also enriched, including BMPR2; FZD2,7, and 1; EGFR; PDGFRB; several Ephrin receptors; ROR2; and DDR2. DDR2 is a receptor tyrosine kinase that binds collagen to initiate signaling and is a transcriptional target of Twist1 in the cranial neural crest (Bildsoe et al., 2016; Vogel et al., 1997). The neuroendocrine specific peptide VGF is induced by the receptor tyrosine kinase BDNF (NTRK2), both of which were present in our data set (Bozdagi et al., 2008). Furthermore, the ligand and receptor pairs enriched in organoid samples in Figure 4 are different from the set of ligands used during the differentiation protocol.

**Figure 4.**
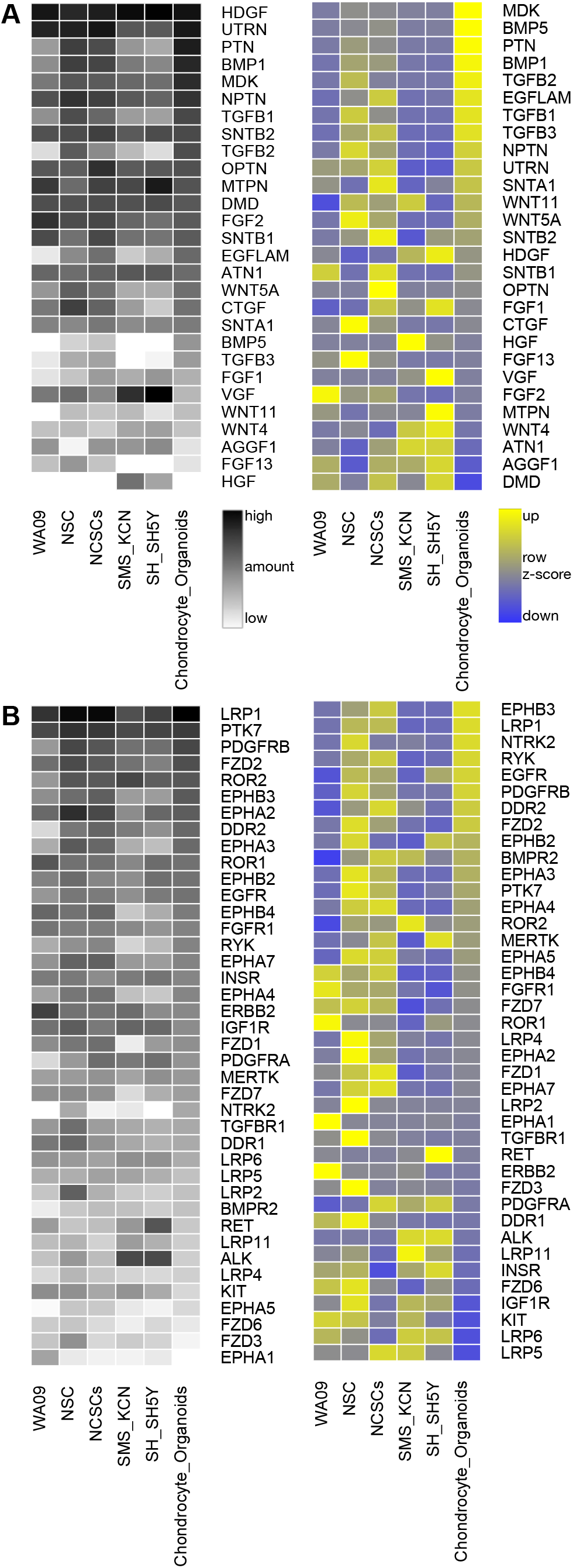
Highlights from mass spectrometry data to examine growth factors (**A**) and receptors (**B**) as markers for differentiation, graphed as in Figure 3. Data are ordered from highest to lowest by the chondrocyte organoid column.

## Discussion

Here we have described the differentiation of craniofacial cartilage organoids from human embryonic stem cells, utilizing a neural crest cell intermediate. Previous studies have demonstrated the differentiation of neural crest lineage chondrocytes (Umeda et al., 2015); our protocol builds from these studies by generating a self-organizing chondrocyte population. Starting with human embryonic stem cells (hESCs), we first differentiated the neural crest lineage from specified neuroectoderm, merging several previously published protocols. Following neural crest isolation, we then differentiated chondrocytes, which self-organized in tissue culture dishes (Figure 1). Cartilage organoids exhibited markers of both cartilage and neural crest cells, indicative of their differentiation from an intermediate lineage (Figure 2).

Previous protocols that have generated articular cartilage for transplantation and tissue regeneration have utilized different approaches. Most cartilage tissue models first derive or obtain mesenchymal lineage cells from patient-derived adult bone marrow stem cells and adipose-derived stem cells, which can then be differentiated into articular cartilage via the application of specific growth factors (Burnsed et al., 2016). This approach can be advantageous because these cells are more differentiated and less likely to form tumors in transplanted tissue. They also start from single-cell suspensions, which are amenable to bioprinting techniques (Leberfinger et al., 2017). However, these cell lineages are not ideal for craniofacial cartilage transplantations. First, for many facial structures, there is no direct underlying bone that can act as a cell seeding scaffold or as printing pads for chondrocytes. Second, craniofacial cartilage is distinct in that it does not have to withstand the large amounts of stress that is applied to articular cartilage. Finally, cartilage is a terminal differentiation stage of neural crest derived chondrocytes, whereas mesenchymal stem cells may be more likely to further ossify into bone. Therefore, protocols that utilize a neural crest cell intermediate to derive craniofacial cartilage are more appropriate for the regeneration of facial structures.

To start the differentiation procedure from an embryonic cell or induced pluripotent cell lineage, three main techniques are typically used to induce mesenchymal cell formation; embryoid body (EB) formation, pellet cell culture, and micromass culture. Suchorska et al. recently compared the efficacy of each of these approaches and found that embryoid body culture was the most efficient and least challenging of the techniques (Suchorska et al., 2017b). Indeed, EB formation appears to be the most commonly used technique to derive mesenchymal stem cells in the laboratory, although these cell lineages often suffer from hypertrophy and osteogenic differentiation (Chen et al., 2015; Pelttari et al., 2006). The protocols described by Suchorska et al. utilized BMP-2, TGF-β1, and TGF-β3 addition to the EB culture media, members of growth factor families that were represented in our approach. However, they also found that addition of growth factors in excess concentration was more likely to induce osteogenesis, and changed the expression of collagen isoforms to a pattern that resembled bone (Suchorska et al., 2017a; Suchorska et al., 2017b). These findings highlight one advantage of our protocol; by forming an autocrine signaling loop, chondrocyte cultures may be less prone to hypertrophy by controlling the amount of growth factors produced or by controlling the response to growth factor induced signaling. Because we did not want mesoderm-derived cartilage, we did not use the embryoid body method to derive chondrocytes. However, our methods followed a similar line of reasoning by mimicking the normal development of the neural crest by first differentiating the neuroectoderm lineage, inducing neural rosette formation, and finally isolating cells that expressed neural crest markers.

To differentiate NCSCs, we combined several previously published protocols (Lee et al., 2010; Leung et al., 2016a; Mica et al., 2013; Zeltner et al., 2014). Because our goal was to derive chondrocytes, the purity of the intermediate NCSC population before sorting was less of a concern. Fluorescence activated cell sorting (FACS) of NCSC cultures provided a more uniform NCSC population, but contributed to high amounts of cell death and severely decreased the NCSC yield. Therefore, we fit the NCSC differentiation to a shorter timeline, and purified NCSCs via magnetic-activated cell sorting (MACS) to increase cell survival.

The mass spectrometry data indicated that cartilage organoids concomitantly express growth factors and receptors for those growth factors, suggesting that an autocrine loop forms to drive chondrocyte differentiation (Figure 4). Both pleiotrophin (PTN) and midkine (MDK), which are heparin-binding growth factors, activate signaling pathways through the receptor tyrosine kinase, ALK and are well known to be involved in chondrogenesis during mouse development and bone repair (Dreyfus et al., 1998; Haffner-Luntzer et al., 2014; Ohta et al., 1999). TGF-β family members are also involved in chondrogenesis, with all three isoforms (TGF-β1,2, and 3) represented in mesenchymal progenitor populations (Wang et al., 2015). In particular, TGF-β1 increases collagen production in embryonic stem cells, while TGF-β2 and -β3 increase glycosaminoglycan production in human bone marrow mesenchymal stem cells (Barry et al., 2001; Yang et al., 2009). There is less evidence in the literature for BMP5 involvement during these processes, especially in human models, and we were surprised to see it more highly represented in our data set compared to the canonical cartilage growth factors BMP2 and BMP4. However, in rats, BMP5 expression has been shown to increase during tibia growth plate development, and inhibition of BMP5 signaling significantly decreased extracellular matrix production in primary rat chondrocytes (Mailhot et al., 2008). In an older study, Kingsley et al. found that mice harboring mutations and deletions in the short ear gene locus also exhibited deletions in the BMP5 coding region, and were a likely cause for the skeletal defects seen in short ear mice (Kingsley et al., 1992; Nie et al., 2006). Guenther et al. then found that specific BMP5 enhancer regions controlled cartilage and bone growth zones induced by BMP5 expression, which included cartilage deposits in the mouse nasal cavity (Guenther et al., 2008). In zebrafish, BMP5 is required for neural crest progenitor survival, and a BMP5 morpholino significantly reduced Alcian-blue cartilage staining, though this could be due to loss of progenitor NC populations and not specifically NC derived chondrocytes (Shih et al., 2017). To our knowledge, a thorough analysis of the contribution of BMP5 signaling to human craniofacial development has not been done, although insights from animal models, and our present study, suggest that BMP5 plays a role in both neural crest differentiation and in cartilage formation.

While several groups have differentiated chondrocytes from stem cells, to our knowledge using derived human iPS cells to generate large amounts of transplantable cartilage has not yet been achieved. Current approaches for facial reconstruction usually involve autologous chondrocyte implantation (ACI) to the damaged area, which involves harvesting patient cartilage from an un-injured location in the face, typically from nasal cartilage (Watson and Reuther, 2014). Nasal septal chondrocytes are neural crest lineage chondrocytes and retain some proliferative potential (Pelttari et al., 2017). For example, isolated human nasal septal chondrocytes have been successfully grafted to mice, expanded, applied to synthetic polymers to facilitate injection, and then used for reconstruction (Dobratz et al., 2009). Furthermore, NCSCs have been identified within isolated dental pulp stem cell (DPSC) populations (Janebodin et al., 2011), which have been successfully used to regenerate mandibular bone (d’Aquino et al., 2007; Gronthos et al., 2000). Isolated DPSCs were limited to bone re-growth, but clonal DPSCs could differentiate into other neural crest derived cell lineages (Janebodin et al., 2011). Together, these studies highlight the potential applications of different NC lineages to craniofacial regeneration and reconstruction.

How different NC lineages colonize and differentiate within facial structures, and how this is affected by different signaling pathways is an open area of research. To further determine the contribution of each signaling pathway to craniofacial development, our future goals involve an analysis comparing the efficiency of organoid growth and the activation of signaling molecules downstream of the ligand and receptor pairs we identified as part of an autocrine signaling loop. The presence of an autocrine signaling loop may be exploited if organoids, or the underlying chondrocyte population, are intended for transplantation or growth on cell scaffolding matrices. However, more work is needed to determine the exact growth factor formulation that most efficiently produces craniofacial cartilage for these types of experiments, and may involve other growth factors identified in our data set. Recent work by Kaucka et al. tracked different neural crest lineages throughout mouse cranial development and found that ectomesenchymal neural crest cells could contribute equally to osteogenic, chondrogenic, odontogenic, and adipogenic lineages within a clonal niche in the developing facial structure (Kaucka et al., 2016). Determining the differentiation potential of neural crest derived chondrocytes, and other cells within organoids, will be critical for experiments exploring organoid growth for transplantations and cartilage repair. Furthermore, addressing these questions in a human context could identify similarities and differences between human and animal models, while shedding light on the mechanisms that influence neural crest differentiation within specific tissue types.

The organoids described here are a promising model for human craniofacial cartilage development (Drubin and Hyman, 2017). Our findings demonstrate that they retain markers of both their neural crest lineage, while also exhibiting the structural and molecular markers of cartilage. Because they migrate and self-organize, they may be amenable to transplantation and growth on cell scaffolding materials. If organoids can be grown into specific shapes or on scaffolding biomaterials, they would be a promising candidate for the regeneration and repair of craniofacial defects. Therefore, these organoids are applicable both to the biology of human craniofacial development and to the improvement of stem cell models for craniofacial regeneration and repair.

## Materials and Methods

Please see the materials table at the end of this section for product details and exact media formulations.

### Cell Culture

WA09 hESCs were purchased from the WiCell Institute. hESCs were expanded and maintained in mTeSR™1 (Stem Cell Technologies), passaged with Versene, and cultured on plates coated with growth factor reduced Matrigel (Corning). All cell lines were grown in a humidified incubator at 37°C, 5% CO2.

### Neural Crest Differentiation

Neural Crest Stem Cells (NCSCs) and Neural Stem Cells (NSCs) were derived by integrating the approaches of several previously published protocols (Chambers et al., 2015; Kreitzer et al., 2013b; Lee et al., 2010; Leung et al., 2016b; Mica et al., 2013; Zeltner et al., 2014). WA09 hESCs were passaged in Matrigel coated dishes to 25-40% confluency prior to differentiation. Daily changes with different media compositions and the addition of growth factors and small molecule inhibitors were used to direct differentiation. Day 1: media was changed from mTeSR1 to KSR containing 0.1 µM LDN193189 and 10 µM SB431542. Day 2: 75% KSR, 25% N2 containing 10 µM SB431542 and 3 µM CHIR99021. Day 3: 50% KSR, 50% N2 containing 3 µM CHIR99021. Day 4: 25% KSR, 75% N2 containing 3 µM CHIR99021. Day 5: 100% N2 containing 3 µM CHIR99021. Day 6-9: media was changed daily with NC Specification media containing 10 µM ROCK inhibitor Y-27632. On day 10 after differentiation, NCSCs were sorted via magnetic cell sorting. p75+ NCSCs were positively selected following the MACS MS protocol (Miltenyi Biotech). p75-flow-through was collected as “NSCs”, although this population represents a multitude of cell types. NCSCs were plated and maintained by passaging on Poly-L-Ornithine/Laminin/Fibronectin coated dishes in NCSC Maintenance media. NSCs were plated and maintained by passaging on Matrigel coated dishes in NSC media. Overly confluent NCSC cultures spontaneously form and grow free-floating cartilage organoids when fed regularly with NCSC maintenance media. Organoids were harvested with a sterile spatula for cryosectioning.

### Organoid Cryosections

Cartilage organoids were fixed in fresh 4% PFA in 1X PBS, rotated at 4°C overnight. Organoids were dehydrated in the following stepwise sucrose washes at 4°C: 5% sucrose - 2 hrs., 10% sucrose - 2 hrs., 30% sucrose - overnight, 1:1 ratio of 30% sucrose:OCT - 2 hrs., 100% OCT - 2 hrs. Organoids were embedded in OCT and snap frozen in a dry ice and ethanol bath. Samples were stored at -80°C before sectioning. Cryosectioning was performed using a Leica CM1950 cryostat. 16 µm thick slices were collected at -25°C on Superfrost plus slides (Fisherbrand). Slices were air dried before storage at -80°C.

### Immunofluorescence

Before staining, frozen slides were dried at room temperature overnight. Slices were rehydrated at room temperature in 1X TBS for 20 minutes, and permeabilized for 45 minutes (Permeabilization solution: 0.1% saponin, 1% BSA, 2% Goat serum in 1X TBS). Primary antibodies were diluted in Permeabilization solution, added to samples, and incubated overnight at 4°C. Slides were washed with Perm. four times. Secondary antibody diluted in Perm. was added to samples and incubated for 1 hour at room temperature. Sections were mounted with SlowFade Gold antifade reagent with DAPI (Invitrogen) and imaged on an Olympus FV1000 confocal microscope.

### Simple Stains

Safranin O staining: Frozen slides were rehydrated in distilled water, stained with Weigerts iron hematoxylin for 10 minutes, washed in running tap water for 10 minutes, stained with 0.05% fast green for 5 minutes, rinsed with 1% acetic acid for 10 seconds, stained in 0.1% safranin for 5 minutes, dehydrated in 95% ethanol for 2 minutes (2X), 100% ethanol for 2 minutes (2X), and finally xylene for 2 minutes (2X).

Toluidine Blue staining: Frozen slides were rehydrated in distilled water, stained in toluidine blue solution (3.5mM in 1% NaCl, 7.7% ethanol, pH 2.38), washed 3X with distilled water, dehydrated: 10 dips in 95% ethanol, 10 dips in 100% ethanol, 10 dips in 100% ethanol, cleared in xylene 2X 3 minutes each, and mounted with mounting media and coverslip media.

Hematoxylin and Eosin: H&E stained slides of organoid cryosections were prepared and provided by Dr. Brad Peterson M.D. (Staff Pathologist, Department of Pathology, Community Medical Center, 2827 Fort Missoula Road, Missoula, Montana 59804).

### Mass Spectrometry Sample Preparation

Cells were washed and harvested in either phosphate-buffered saline or versene. Organoids were harvested with a sterile spatula. Cell pellets/organoids were stored at -80°C. Cells were lysed in a 10:1 (v/w) volume of lysis buffer [5% SDS, 100 mM NaCl, 20 mM Hepes (pH 8.5), 5 mM dithiothreitol, 2.5 mM sodium pyrophosphate, 1 mM β-glycerophosphate, 1 mM Na3VO4, and leupeptin (1 mg/ml)], and proteins were reduced at 60°C for 30 min. Proteins were then alkylated by the addition of 10 mM iodoacetamide (Sigma-Aldrich) for 45 minutes at room temperature in the dark, and methanol/chloroform precipitation was performed. Protein pellets were resuspended in urea lysis buffer (8 M urea, 20 mM Hepes (pH 8.5), 1 mM sodium orthovanadate, 2.5 mM sodium pyrophosphate, and 1 mM β-glycerolphosphate) and sonicated. Cell lysates were diluted to 2 M urea in 20 mM Hepes (pH 8.5) and 1mM CaCl2 for Lys-C digestion overnight at 37°C, then diluted twofold followed by trypsin (Promega) digestion for 4 hours at 37°C. Samples were then acidified to pH 2 to 3 with formic acid, and peptides were purified on a Waters Sep-Pak column and dried in a speed-vac. Peptides were quantified using a micro-BCA assay (ThermoFisher). Peptides were crosslinked to ten mass tag labels (ThermoFisher TMTplex) and analyzed on an Orbitrap Fusion Lumos (ThermoFisher; at Cell Signaling Technology, Beverly, MA). Identification of peptides and quantification of mass tags was obtained from the MS2 spectrum after fragmentation by MS/MS analysis as described (Beausoleil et al., 2006; Grimes et al., 2018; Guo et al., 2014; Possemato et al., 2017; Stokes et al., 2016). The iBAQ method was used to normalize signals, where a proteins total intensity is divided by the number of tryptic peptides between 6 and 30 amino acids in length (Arike et al., 2012).

**Table 1:**
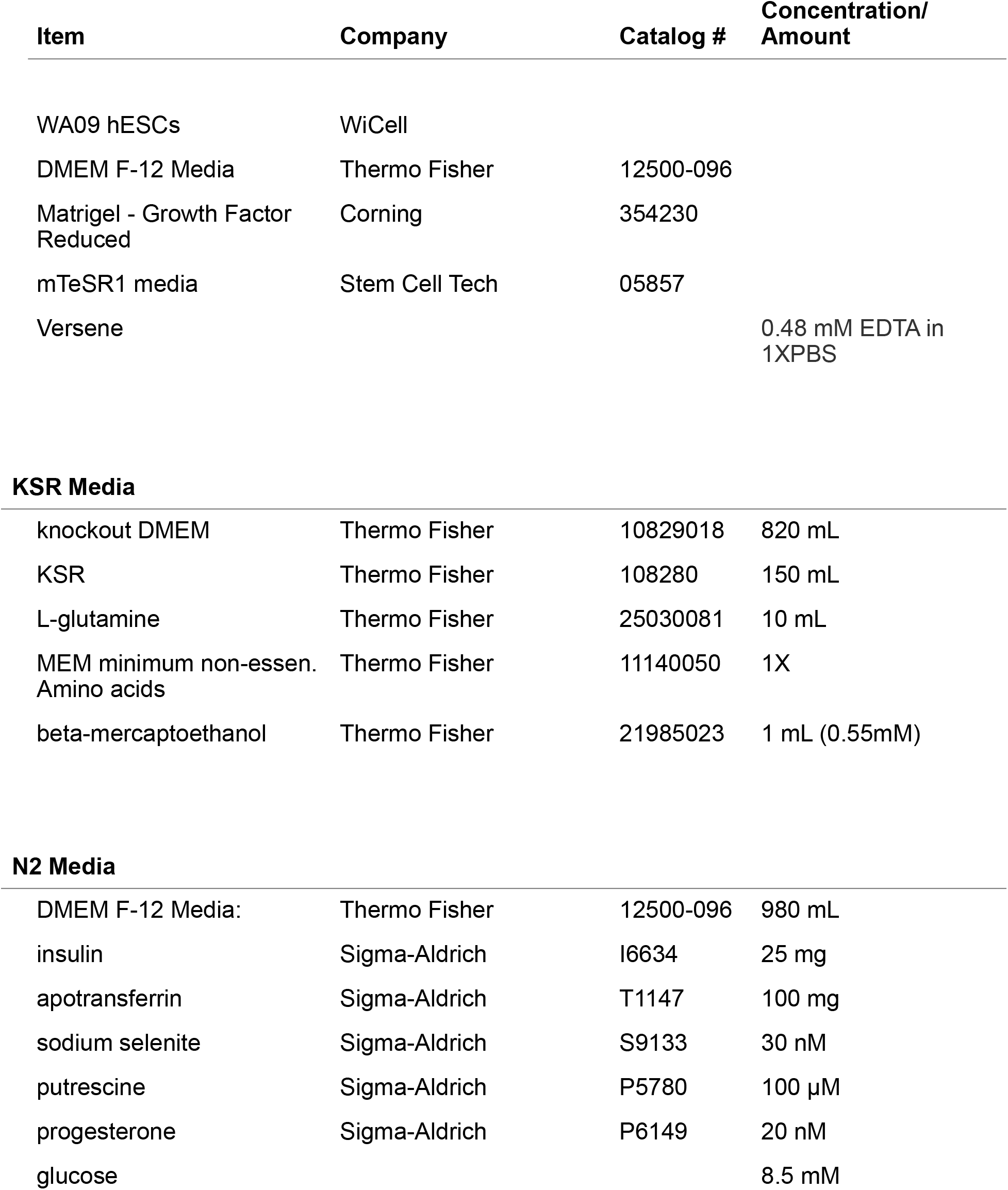
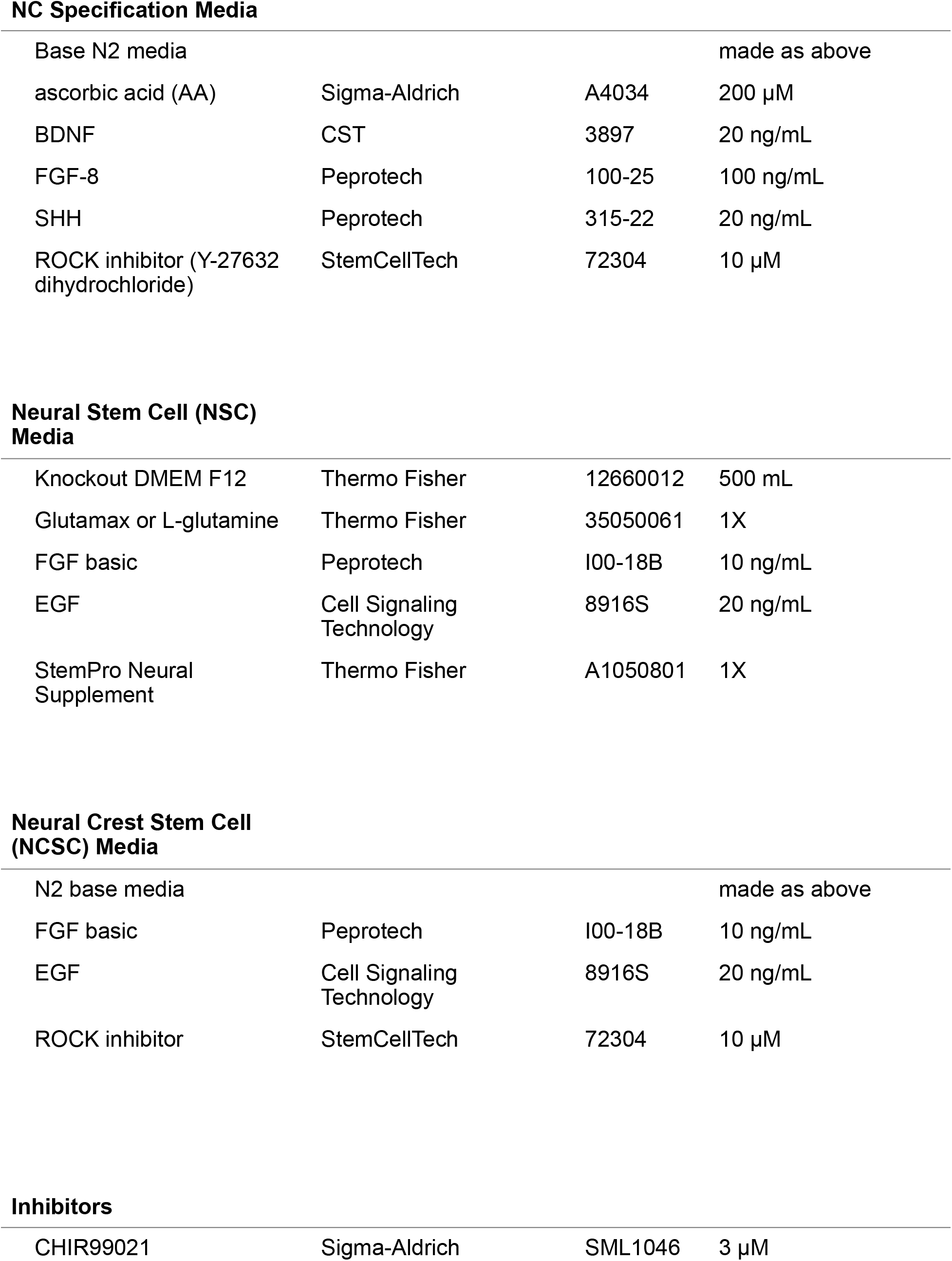
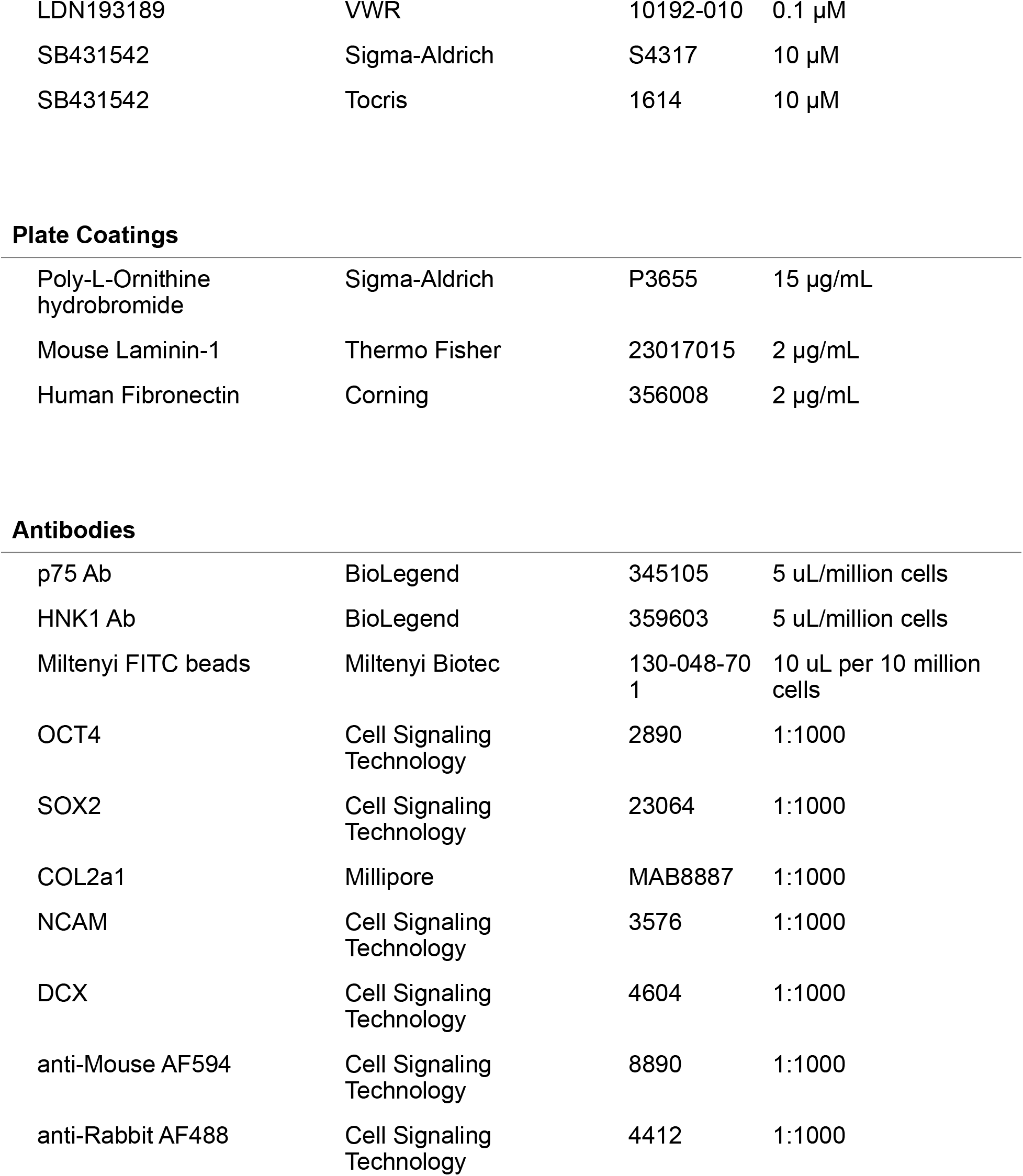
Media and reagents.

| Item | Company | Catalog # | Concentration/<br>Amount |
| --- | --- | --- | --- |
| WA09 hESCs | WiCell |  |  |
| DMEM F-12 Media | Thermo Fisher | 12500-096 |  |
| Matrigel - Growth Factor Reduced | Corning | 354230 |  |
| mTeSR1 media | Stem Cell Tech | 05857 |  |
| Versene |  |  | 0.48 mM EDTA in 1XPBS |
| **KSR Media** |  |  |  |
| knockout DMEM | Thermo Fisher | 10829018 | 820 mL |
| KSR | Thermo Fisher | 108280 | 150 mL |
| L-glutamine | Thermo Fisher | 25030081 | 10 mL |
| MEM minimum non-essen. Amino acids | Thermo Fisher | 11140050 | 1X |
| beta-mercaptoethanol | Thermo Fisher | 21985023 | 1 mL (0.55mM) |
| **N2 Media** |  |  |  |
| DMEM F-12 Media: | Thermo Fisher | 12500-096 | 980 mL |
| insulin | Sigma-Aldrich | I6634 | 25 mg |
| apotransferrin | Sigma-Aldrich | T1147 | 100 mg |
| sodium selenite | Sigma-Aldrich | S9133 | 30 nM |
| putrescine | Sigma-Aldrich | P5780 | 100 µM |
| progesterone | Sigma-Aldrich | P6149 | 20 nM |
| glucose |  |  | 8.5 mM |
| **NC Specification Media** |  |  |  |
| Base N2 media |  |  | made as above |
| ascorbic acid (AA) | Sigma-Aldrich | A4034 | 200 μM |
| BDNF | CST | 3897 | 20 ng/mL |
| FGF-8 | Peprotech | 100-25 | 100 ng/mL |
| SHH | Peprotech | 315-22 | 20 ng/mL |
| ROCK inhibitor (Y-27632 dihydrochloride) | StemCellTech | 72304 | 10 μM |
| **Neural Stem Cell (NSC) Media** |  |  |  |
| Knockout DMEM F12 | Thermo Fisher | 12660012 | 500 mL |
| Glutamax or L-glutamine | Thermo Fisher | 35050061 | 1X |
| FGF basic | Peprotech | 100-18B | 10 ng/mL |
| EGF | Cell Signaling Technology | 8916S | 20 ng/mL |
| StemPro Neural Supplement | Thermo Fisher | A1050801 | 1X |
| **Neural Crest Stem Cell (NCSC) Media** |  |  |  |
| N2 base media |  |  | made as above |
| FGF basic | Peprotech | 100-18B | 10 ng/mL |
| EGF | Cell Signaling Technology | 8916S | 20 ng/mL |
| ROCK inhibitor | StemCellTech | 72304 | 10 μM |
| **Inhibitors** |  |  |  |
| CHIR99021 | Sigma-Aldrich | SML1046 | 3 μM |
| LDN193189 | VWR | 10192-010 | 0.1 µM |
| SB431542 | Sigma-Aldrich | S4317 | 10 µM |
| SB431542 | Tocris | 1614 | 10 µM |
| Plate Coatings |  |  |  |
| Poly-L-Ornithine hydrobromide | Sigma-Aldrich | P3655 | 15 µg/mL |
| Mouse Laminin-1 | Thermo Fisher | 23017015 | 2 µg/mL |
| Human Fibronectin | Corning | 356008 | 2 µg/mL |
| Antibodies |  |  |  |
| p75 Ab | BioLegend | 345105 | 5 uL/million cells |
| HNK1 Ab | BioLegend | 359603 | 5 uL/million cells |
| Miltenyi FITC beads | Miltenyi Biotec | 130-048-701 | 10 uL per 10 million cells |
| OCT4 | Cell Signaling Technology | 2890 | 1:1000 |
| SOX2 | Cell Signaling Technology | 23064 | 1:1000 |
| COL2a1 | Millipore | MAB8887 | 1:1000 |
| NCAM | Cell Signaling Technology | 3576 | 1:1000 |
| DCX | Cell Signaling Technology | 4604 | 1:1000 |
| anti-Mouse AF594 | Cell Signaling Technology | 8890 | 1:1000 |
| anti-Rabbit AF488 | Cell Signaling Technology | 4412 | 1:1000 |

